# Evaluation of Antigen Expression and Early Immune Response following Cutaneous Suction-mediated DNA Delivery

**DOI:** 10.1101/2025.09.15.676388

**Authors:** Nandita C. Jhumur, Yukyung Um, Sarah H. Park, Emran O. Lallow, Christine C. Roberts, Jerry W. Shan, Jonathan P. Singer, Jeffrey D. Zahn, Young K. Park, Lisa K. Denzin, David I. Shreiber, Joel N. Maslow, Hao Lin

## Abstract

Suction-based in vivo cutaneous DNA transfection is a newly developed, cost-effective method that produces high transfection efficiency. This method has shown robust immunogenic responses following SARS-CoV-2 DNA vaccination in both pre-clinical studies and clinical trials. The current work investigates suction-based transfection and immune activation on a detailed, cellular level. The spatiotemporal patterns of antigen expression in rat skin following suction-induced delivery of a pEGFP-N1 plasmid and a SARS-CoV-2 DNA vaccine are evaluated via immunofluorescence staining, which demonstrates early and prolonged expression. The epidermis is identified as the primary location of transfection, and the transfected cells are primarily epidermal keratinocytes. Early immune response is assessed by detection of antigen presenting cells (APCs) following suction-induced DNA vaccination.

## 1. Introduction

In vivo DNA transfection has broad applications in numerous fields including DNA vaccination, gene therapy, cancer and stem cell research, among many others.^[1–4]^ Although chemical and biological means of transfection have been employed in vivo, physical methods are now the predominant approach, which includes delivery via electroporation, gene gun, jet injection, sonoporation, optical transfection, hydrodynamic transfection, and magnetofection, among others.^[5–16]^ While physical transfection methods avoid potential adverse effects associated with chemical or biological agents,^[17, 18]^ they commonly cause pain and/or tissue damage or injury.^[19]^

The skin is an accessible, advantageous location for in vivo DNA delivery. Additionally, the skin tissue is rich in antigen presenting cells (APCs) including Langerhans cells (LCs) and dendritic cells (DCs) that circulate through the skin to take up foreign antigens, carry them to local lymph nodes, and trigger adaptive immune response following vaccination.^[20, 21]^ We have previously developed a suction-based in vivo DNA transfection method where a negative pressure is applied to the skin surface following intradermal (ID) injection (Figure 1).^[22]^ A systematic study of key mechanical parameters revealed that antigen expression is strongly correlated to patterns of tissue strain and tissue tension.^[17]^ Suction-based delivery method demonstrated a strong immune response to a SARS-CoV-2 DNA vaccine in both pre-clinical studies and clinical trials (NCT04673149; NCT05182567).^[23–25]^ Compared with other physical DNA vaccination methods, such as jet injection and electroporation, suction-based cutaneous delivery resulted in similar humoral and higher cellular immune responses.^[24, 26]^

**Figure 1:**
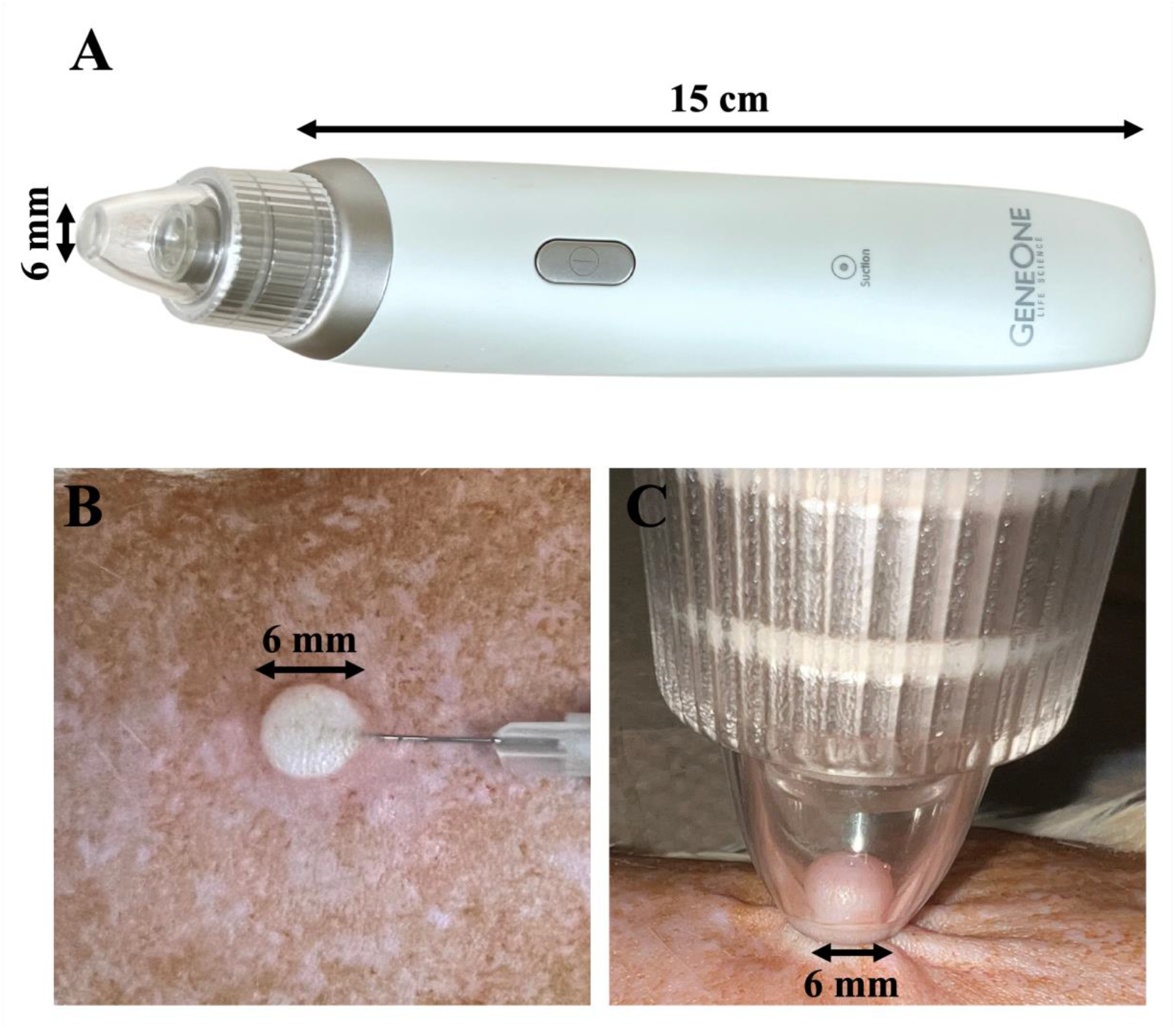
(A) Suction-application Device. (B) 50 µL ID injection in rat skin using the Mantoux technique. (C) Application of the suction device on rat skin post-ID injection.

In this work, we investigate the spatiotemporal patterns of cutaneous suction-mediated DNA transfection at a cellular level. Transgene expression of GFP and the SARS-CoV-2 spike S1 protein delivered via plasmids was determined by immunofluorescence and histological analysis to examine the location and cell types involved in antigen expression, as well as to detect the influx of antigen presenting cells (APCs) as the initial step in immunogenesis.

## 2. Materials and Methods

### 2.1. Animals

Murine-pathogen free, standard male Sprague-Dawley rats (NTac-SD, 7-11 weeks old) were purchased from Taconic Biosciences, Inc. (Germantown, NY). The rats were kept in a controlled environment according to Rutgers University Institutional Animal Care and Use Committee (IACUC), protocol: 201800077.

### 2.2 DNA plasmids

The GLS-5310 SARS-CoV-2 DNA vaccine and the GLS-5300 MERS-CoV DNA vaccine were provided by GeneOne Life Science, Inc. (Seoul, Republic of Korea). Both vaccines were manufactured under cGMP conditions at VGXI, Inc. (Conroe, TX). Pre-clinical grade pEGFP-N1 plasmid DNA was purchased from GenScript (Piscataway, NJ). For the GFP transfection time study (Figure 2), 25 μg pEGFP-N1 plasmid was prepared in a 50 μL solution using 1× PBS. For the rest of the studies, a cocktail of 25 μg pEGFP-N1 plasmid and 150 μg SARS-CoV-2 DNA vaccine (GLS-5310) was prepared in a 50 μL solution using an SSC formulating buffer (GLS-0001) provided by GeneOne.^[26]^ A 1μg/μL solution of MERS-CoV DNA vaccine (GLS-5300) was prepared in 50 μL PBS. Each injection was carried out intradermally into rat dorsal skin with a 28G insulin syringe (BD, Franklin Lakes, NJ) using the Mantoux injection technique.

**Figure 2:**
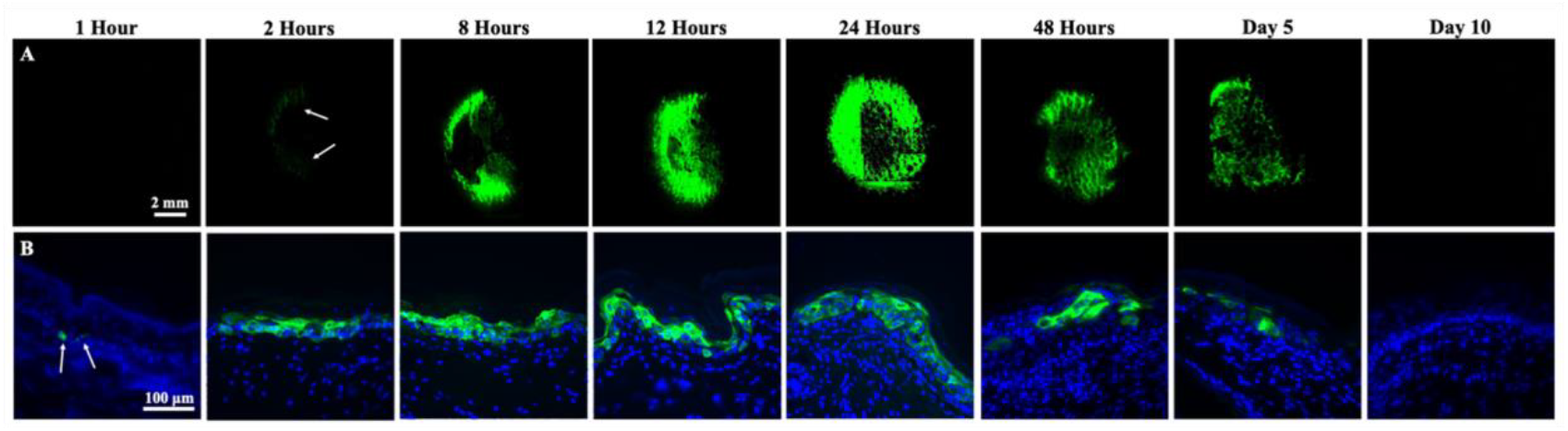
(A) Immunofluorescent images of GFP expression (top view) in rat skin. (B) Immunolabeled GFP in rat skin cross-section (thickness: 10 µm) at various time points up to 10 days following suction-induced pEGFP-N1 transfection.

### 2.3 Suction application

The GeneDerm device was provided by GeneOne and used for the application of suction (Figure 1A) with the following specifications: nozzle inner diameter, 6 mm; suction pressure, 80 kPa; suction application time, 30 seconds. The rat dorsal skin was prepared for injection as described in our prior work.^[17]^ The diameter of each 50 μL ID injection bleb was approximately 6 mm (Figure 1B). Suction was performed on the injection bleb using the GeneDerm device immediately after injection (Figure 1C). All the rats were housed individually and were euthanized via CO_2_ asphyxiation at one of the time points for ensuing analysis: 1 hour, 2 hours, 6 hours, 8 hours, 12 hours, 18 hours, 24 hours, 48 hours, day 5, or day 10.

### 2.4 Tissue processing, immunofluorescence staining, and imaging

The treated area of the rat skin was dissected at the time points indicated above. The excised skin blocks were placed on glass slides (Colorfrost Plus Microscope slides, Fisher Scientific, Waltham, MA) and imaged with an Olympus IX81 inverted epifluorescence microscope (Olympus, Tokyo, Japan) to detect GFP expression. The skin blocks were then fixed in 10% Neutral Buffered Formalin (G-Biosciences, St. Louis, MO) for 24 hours. The fixed tissue blocks were incubated in 10% sucrose-PBS solution overnight at 4 °C. Afterward, they were embedded into the Tissue-Tek O.C.T. Compound (Sakura Finetek, Torrance, CA) and stored in a −80°C freezer. The frozen tissues were cryo-sectioned into 10 μm thick skin cross-sections by the Rutgers Research Pathology Services and mounted onto electrostatically surface-treated glass slides (Colorfrost Plus Microscope slides, Fisher Scientific, Waltham, MA). Tissue sections on slides were then washed in tris-buffered saline with 0.1% Tween 20 detergent (TBST) for 5 minutes three times. Next, they were incubated in a permeabilization and blocking buffer (1% BSA, 0.25% Casein, 0.1% Cold fish skin gelatin, TBS, pH-7.4, with 1.0% Tween 20, 0.3% Triton X-100, and 150 mM Sodium azide) with 5% Normal Goat Serum (Thermo Fisher Scientific Inc., Waltham, MA) for 30 minutes to block non-specific binding and then incubated in a solution of the primary antibodies diluted in an incubation buffer (1% BSA in TBS, pH-7.4 with 0.1% Tween 20, 0.015 mol/L Sodium azide) overnight at 4 °C. The sections were washed in TBST for 5 minutes three times and incubated for 30 minutes in a secondary antibody solution. The slides were again washed in TBST for 5 minutes three times and cover-slipped with ProLong Diamond Antifade Mountant with DAPI (Thermo Fisher Scientific Inc., Waltham, MA). An Olympus IX81 inverted epifluorescence microscope (Olympus, Tokyo, Japan) was used for imaging all the tissue sections.

The primary antibodies used were: (1) Rabbit anti-GFP Antibody (A-11122, Thermo Fisher Scientific Inc., Waltham, MA), 4 µg/mL for GFP detection; (2) Rabbit anti-SARS-CoV-2 Spike S1 Antibody (GTX135356, GeneTex, Irvine, CA), 1 µg/mL for SARS-CoV-2 spike S1 protein detection; (3) Rabbit anti-SARS-CoV-2 Spike S2/S2’ Antibody (GTX135386, GeneTex, Irvine, CA), 20 µg/mL for SARSCoV-2 spike S2 protein detection; (4) Mouse anti-Human/Rat pan Cytokeratin Antibody (188-10844-1, RayBiotech Life Inc., Peachtree Corners, GA), 4 µg/mL for cytokeratin detection that can detect both acidic (Type I) and basic (Type II) cytokeratins including Keratins 1, 3-6, 8, 10, 14-16, and 19; (5) Mouse anti-Rat RT1B Antibody (205301, BioLegend Inc., San Diego, CA), 20 µg/mL for antigen presenting cell (APC) detection. The secondary antibodies used were: Goat anti-Rabbit IgG (H+L) Cross-Adsorbed Secondary Antibody, Alexa Fluor 546 (A-11010, Thermo Fisher Scientific Inc., Waltham, MA), 4 µg/mL for GFP, SARS-CoV-2 spike S1 protein, and SARS-CoV-2 spike S2 protein detection; Goat anti-Mouse IgG (H+L) Cross-Adsorbed Secondary Antibody, Alexa Fluor 647 (A-21235, Thermo Fisher Scientific Inc., Waltham, MA), 2 µg/mL for cytokeratin and APC detection.

For hematoxylin and eosin (H&E) staining of non-treated rat skin, the excised skin blocks were fixed, cryo-sectioned into 40 µm thick cross-sections, stained with H&E, and imaged by the Rutgers Research Pathology Services according to their standard protocols.

### 2.5 Immunostaining validation

Normal Mouse IgG (NI03, MilliporeSigma, Burlington, MA) was used as isotype control for validating the immunostaining of cytokeratin and APCs at 4 µg/mL and 20 µg/mL, respectively. Rabbit IgG (I5006, MilliporeSigma, Burlington, MA) was used as isotype control for validating the immunostaining of GFP, SARS-CoV-2 spike S1 protein, and SARS-CoV-2 spike S2 protein at 4 µg/mL, 1 µg/mL, 20 µg/ml, respectively.

Immunostaining of both SARS-CoV-2 spike S1 and S2 proteins was validated by comparing with the following control cases (Figure S1): (i) application of SARS-CoV-2 spike S1 and S2 antibodies to non-treated rat skin sections, (ii) application of SARS-CoV-2 spike S1 and S2 antibodies to MERS-CoV vaccinated rat skin sections to ensure the specificity of the antibodies, and (iii) Rabbit IgG (isotype control) application to SARS-CoV-2 vaccinated rat skin.

## 3. Results

### 3.1 Cutaneous suction mediates early and persistent GFP expression in the skin

We extended our prior work that examined GFP expression through 48 hours ^[22]^ to 10 days post treatment. Skin explants were collected at 1, 2, 8, 12, 24, 48 hours and days 5 and 10, with 3-5 treated skin explants analyzed at each time-point. The dissected skin blocks were imaged with epifluorescence microscope using a 4× objective.

As shown in Figure 2A, GFP expression starts to appear at 1-2 hours, gradually increases at 8 hours and 12 hours, and reaches maximal expression at 24 hours. GFP expression is continually detected through day 5. At day 10, no visible GFP expression is detected.

We additionally processed the skin blocks for immunohistochemical analysis and performed immunofluorescence staining to detect GFP (Figure 2B). Rabbit IgG was used as an isotype control to validate the GFP staining, at the same concentration as the applied Rabbit anti-GFP Antibody along with Goat anti-Rabbit IgG (H+L) Cross-Adsorbed Secondary Antibody, Alexa Fluor 546 (not shown). We selected skin cross-sections from high GFP expressing regions of the skin blocks. At 1-hour post-treatment, a few GFP positive cells were detected in the skin cross-sections. As early as 2 hours after treatment, we see a significant increase in the number of GFP positive cells. Unlike the top view, where GFP expression continues to increase until 24 hours, GFP expression in cross-sections is similar at 8, 12, and 24 hours. For both the top view and cross-sections, GFP expression slowly wanes after 24 hours and completely disappears at day 10.

### 3.2 Temporal expression of the SARS-CoV-2 spike S1 protein following suction-mediated transfection

We next assessed the temporal expression pattern of the SARS-CoV-2 spike S1 protein antigen following in vivo suction-mediated transfection of the GLS-5310 DNA vaccine. To facilitate locating GLS-5310 transfected cells, we injected a cocktail of pEGFP-N1 plasmid and SARS-CoV-2 DNA vaccine (25 μg of pEGFP-N1 and 150 μg of GLS-5310 in a 50 μL solution; see Materials and Methods) into rat dorsal skin and applied the same suction of 80 kPa for 30 seconds. Three rats were used for each time-point evaluation with one treatment spot per rat. We first imaged the dissected skin blocks under the epifluorescence microscope for GFP expression to locate the regions of interest for sectioning; immunofluorescence staining was then performed to detect the spike S1 protein.

The results are shown in Figure 3, which largely corroborate with trends observed with GFP: S1 expression was detected as soon as 1-hour post-treatment, increased over time, with peak expression observed at 24 hours, and then declined. Notably, S1 expression remained detectable through day 10.

**Figure 3:**
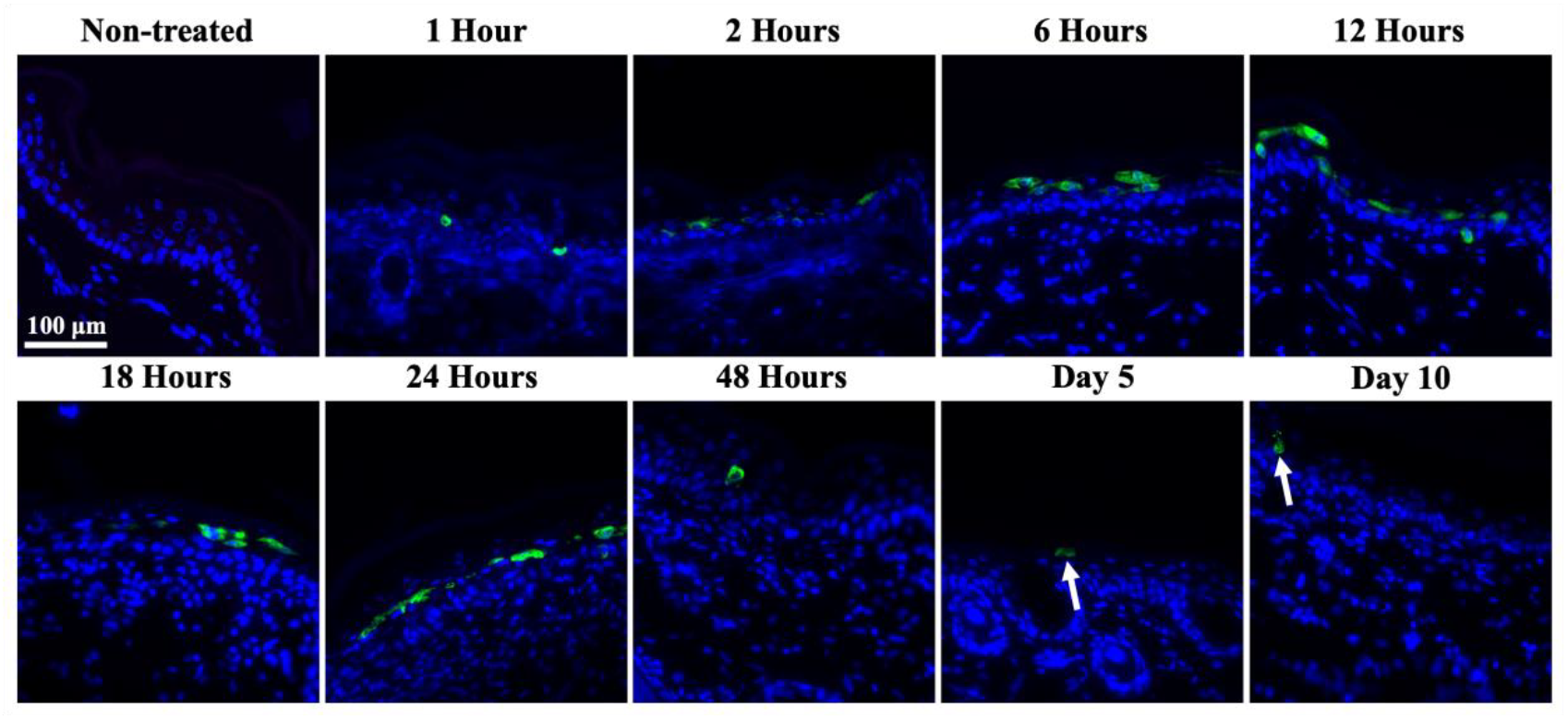
Immuno-labeled SARS-CoV-2 spike S1 protein in rat skin cross-section (thickness: 10 µm) at 1, 2, 6, 12, 18, 24, and 48 hours, and day 5 and 10 following suction-mediated DNA vaccination.

### 3.3. The epidermis is the primary location for suction-induced DNA transfection

The above results indicate that transgene expression is restricted to the superficial layers of the skin. We then further evaluated spatial expression with respect to the tissue structure. A representative H&E-stained skin section in Figure 4A shows different layers of rat skin. The epidermis layer of the skin includes stratum basale, stratum spinosum, stratum granulosum, and stratum corneum.^[27,28]^ Note that the panniculus carnosus layer (robust to rodents but only vestigial to humans) is identified between the fat layer and the deep fascia.^[29]^ In the immunostained images, the different skin layers are identified by the shape and density of the cells.

**Figure 4:**
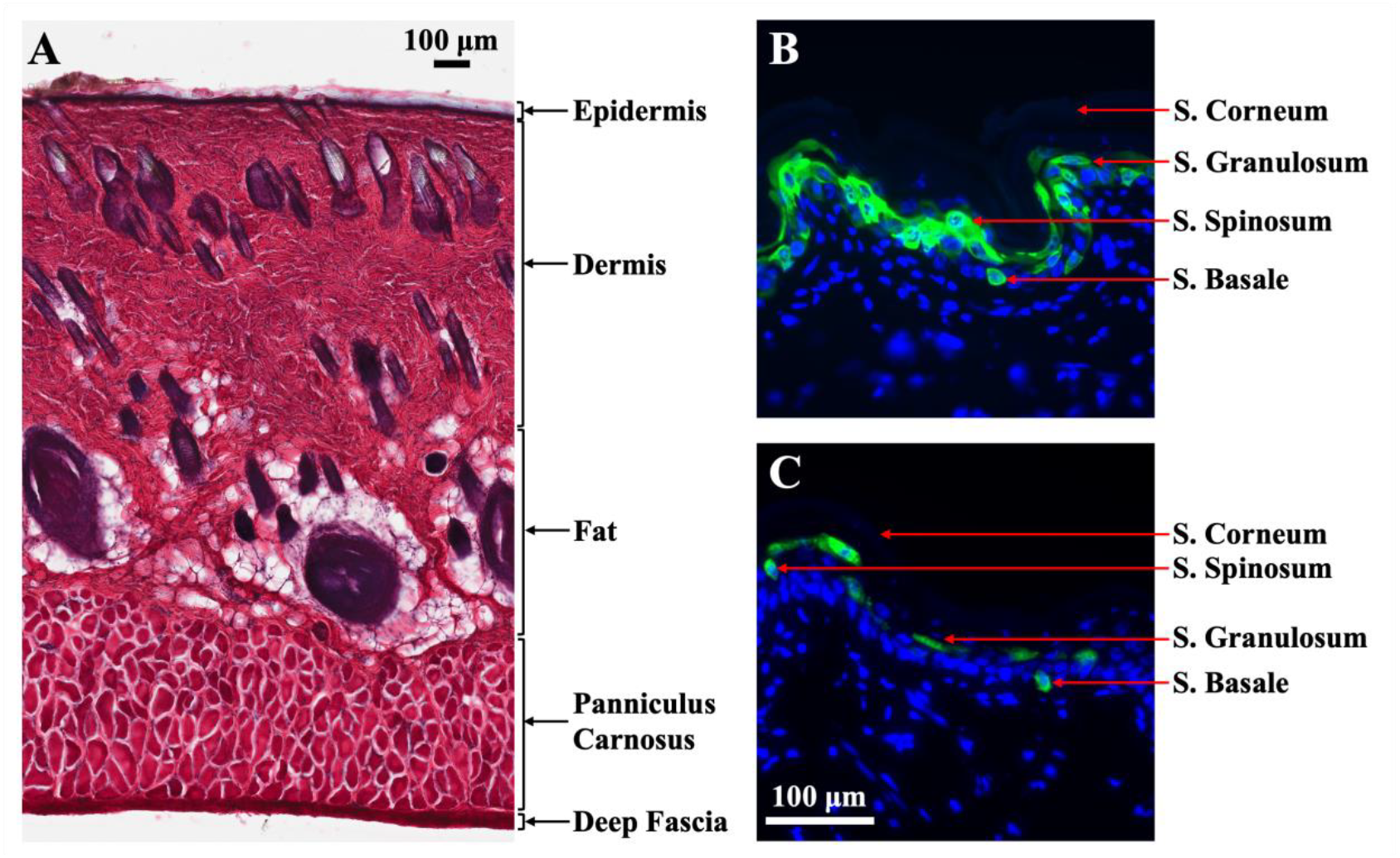
(A) H&E stained image of rat skin cross-section (thickness: 40 µm). (B) Immunostained GFP in rat skin epidermis (section thickness: 10 µm) at 12 hours. (C) Immunostained SARS-CoV-2 spike S1 protein in rat skin epidermis (section thickness: 10 µm) at 12 hours.

**Figure 5:**
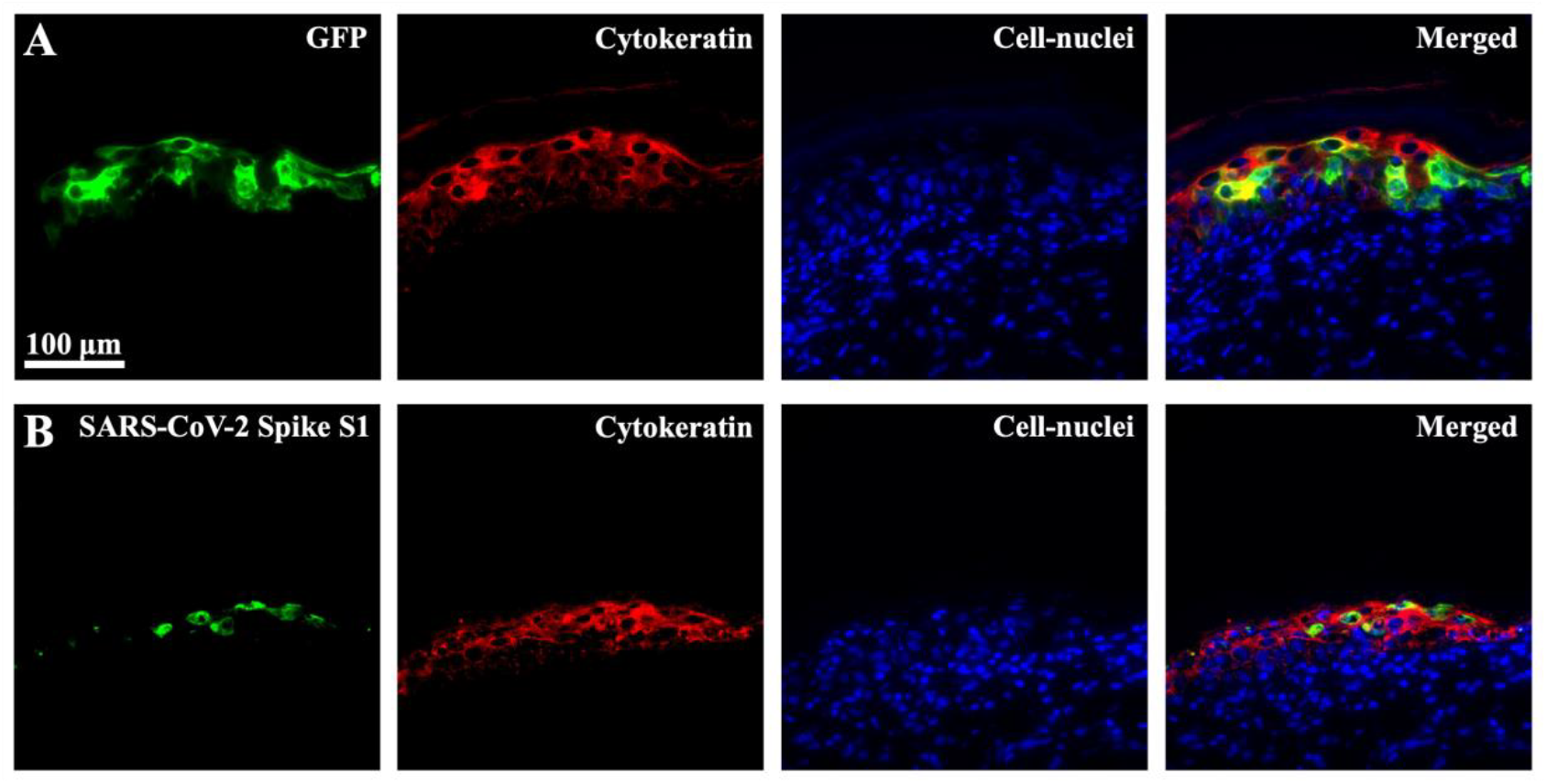
Co-localization of (A) immuno-labeled GFP and cytokeratin and (B) immuno-labeled SARS-CoV-2 spike S1 protein and cytokeratin in rat skin cross-section (thickness: 10 µm) at 24 hours following suction-mediated DNA transfection.

Figure 4 panel B and C show representative images of GFP and S1 protein distribution across the skin epidermis at 12 hours, which is a representative time point of a stable period of expression from Figure 2 and Figure 3, respectively. Figures 2–4 together suggest that the protein expression followed by suction-induced DNA delivery is mainly found in the epidermis. Furthermore, GFP positive cells are located in the stratum basale layer at 1 hour, and populate all three sub-layers of the epidermis including stratum basale, stratum spinosum, and stratum granulosum from 2 hours until day 5 (Figure 2); no protein is detected in the stratum corneum layer since it consists of dead cells. Additionally, no expression has been observed outside of the epidermis. A similar spatial distribution of antigen expression is also observed for the SARS-CoV-2 spike S1 protein. This pattern of expression is also observed with the SARS-CoV-2 spike S2 protein (Figure S1).

### 3.4. Transfected cells are primarily keratinocytes

We next identified the types of cells that were transfected. Due to localization of the transfected cell within the epidermis, we hypothesized that keratinocytes, which are the predominant epidermal cell type, were the main site for transfection.^[27]^

Three rats were injected with the cocktail of pEGFP-N1 and SARS-CoV-2 vaccine in only one site after which suction was applied. The rats were euthanized at 24 hours and the treated skin sites were dissected. Following the selection of representative sections using GFP expression, sections were immunolabeled with either Rabbit anti-GFP or Rabbit anti-spike S1 (SARS-CoV-2) together with Mouse anti-pan cytokeratin antibodies. Mouse IgG was used as an isotype control to validate the immunostaining of Mouse anti-pan cytokeratin (not shown), where no cytokeratin was detected. The merged images show direct co-localization of both the GFP and the spike S1 protein with cytokeratin. Together, these results provide evidence that keratinocytes are the site for suction-mediated transfection.

### 3.5 Antigen presenting cells (APCs) are detected around transfected cells following suction-induced DNA transfection

We examined the early immune response triggered by suction-mediated DNA delivery in rat skin. Three rats were intradermally injected by the same pEGFP-N1 and SARS-CoV-2 DNA vaccine cocktail, followed by suction application, with one site treated per rat. We euthanized the rats at 24 hours and dissected the treated spots for cryo-sectioning and immunofluorescence staining.

We used an anti-rat RT1B antibody to detect the antigen presenting cells (APCs) in rat skin via immunostaining for MHC (Major Histocompatibility Complex) class II molecules. In Figure 6A, representative results from non-treated rats demonstrate low frequency of APCs in the control skin sections. In Figure 6B, no positive immunolabeling was observed in sections of treated skin following immunostaining with Mouse IgG isotype control at the same concentration as the Mouse anti-rat RT1B antibody. Results of co-immunostaining for MHC class II molecules and GFP or spike S1 protein are shown in Figure 6C and 6D, respectively. APCs were detected in all the layers of skin, and similar to the results above, GFP and S1 expression was restricted to the epidermal layers. In Figure 6C and 6D, the merged images show GFP or S1 protein-expressing cells in close proximity to APCs. In general, we found that the overall number of APCs in the treated skin sections is higher compared to the non-treated skin sections, although we did not perform a systematic quantification. Additionally, within the individual APCs, the fluorescence intensity as resulted from immunostaining seems to be higher and more widespread in treated skin sections than that in the non-treated counterpart. Taken together, our results support APC recruitment and/or activation in the epidermal layer in response to antigen production after vaccination via suction, with several of the MHC class II positive APCs directly adjacent to cells expressing GFP or S1.

**Figure 6:**
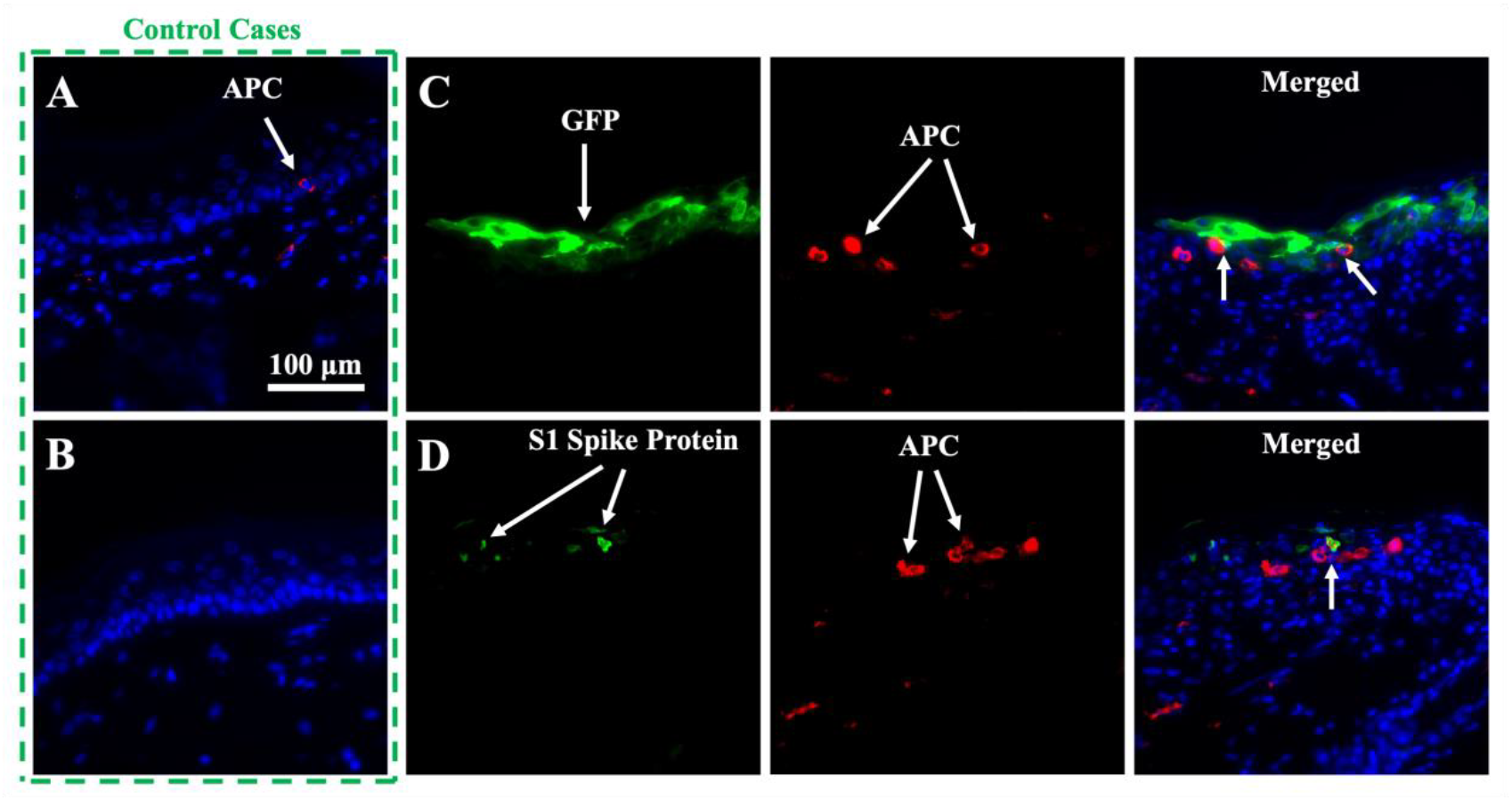
(A) APCs in non-treated rat skin cross-section. (B) Application of Mouse IgG in rat skin cross-section at 24 hours following treatment. (C) Detection of immuno-labeled GFP and APCs in rat skin cross-section at 24 hours following treatment. (D) Detection of immuno-labeled spike S1 protein and APCs in rat skin cross-section at 24 hours following treatment. The skin cross-section thickness is 10 µm for all cases.

## 4. Discussion

Cutaneous suction is a simple-to-implement technique for in vivo DNA delivery and transfection. In our previous work, we optimized the physical parameters and identified major mechanical factors behind the high transfection efficiency by suction application.^[17]^ In this work, we investigated aspects of the transfection process and also examined the initial immune response.

Extending our prior work,^[22]^ we confirm that following suction mediated in vivo transfection, transgene expression is maximal at 24 hours. We then showed that GFP expression is detectable through to 5 days post-treatment, and that S1 expression from the GLS-5310 SARS-CoV-2 DNA vaccine was detected, albeit at low levels, through 10 days post-treatment. Some of the possible causes for this decay include the degradation of the plasmids, cellular turnover, and the immune response triggered by the transfection that clears or transports the antigen.^[30–32]^

GFP and SARS-CoV-2 spike S1 and spike S2 proteins were identified in the epidermis layer of the skin. At 1-hour post-transfection, cells expressing GFP or SARS-CoV-2 spike S1 protein were only found in the stratum basale layer. At later time points, we detected the transfected cells in all three sub-layers of the epidermis including stratum basale, stratum spinosum, and stratum granulosum. From the current results, it is not yet clear if transfection occurs preferentially at the stratum basale. The earlier expression in the stratum basale may be due to the proximity to the injection bleb and thus higher local concentrations of DNA. It may also be correlated with the active proliferation and differentiation process of the basale layer cells.

We noticed that at each time point and when compared with the spike S1 spike protein positive cells, GFP positive cells appeared denser across the epidermis. We hypothesize that this might be due to the higher expression efficiency of the pEGFP-N1 plasmid. The pEGFP-N1 plasmid vector was designed to yield high levels of expression that diffused widely throughout the epidermis. Alternatively, the stability of GFP might be greater than S1, leading to high levels of detectable protein.

The epidermal layer contains various cell types including keratinocytes, melanocytes, Langerhans cells, and Merkel cells. We used an anti-pan cytokeratin antibody to broadly detect the various keratins that are expressed by the keratinocytes in the different sublayers of epidermis.^[33, 34]^ The co-localization of cytokeratin with GFP and with SARS-CoV-2 spike S1 protein in the different sublayers of epidermis suggest that transfected cells are keratinocytes.

The underlying mechanism behind the strong immune response in suction-driven DNA transfection as observed in our prior animal and human studies is of key interest. We used immunolabeling to investigate the early immune response, which might include the recruitment of APCs, likely dermal dendritic cells, to the treated skin site following the suction-induced cutaneous DNA delivery. The main APCs in skin include Langerhans cells and dermal dendritic cells. These cells capture antigens in the skin via phagocytosis or endocytosis and ultimately digest the antigens into peptides for loading onto MHC class I and II molecules. The APCs then traffic to draining lymph nodes where they function to initiate T cell activation and the adaptive immune response.^[35, 36]^ We used an antibody specific for MHC class II to identify APCs in the rat skin (Figure 6). Langerhans cells (LCs), and dermal dendritic cells are the most likely detected cell types, although they cannot be definitively identified using the current immunolabeling approach. We observed an increase in the number of MHC class II positive cells and more intense MHC class II staining in the vaccinated skin. This indicates that an immune response is initiated following the suction-driven DNA transfection.

In summary, we have studied the suction-based DNA transfection at the cellular level, evaluated the location of transfection across skin layers, detected the type of transfected cells, and identified antigen bearing, MHC class II positive APCs in rat skin tissue by immunofluorescence staining. This study improves our understanding of this newly invented technical platform for nucleic acid base biologics delivery. Future mechanistic studies are planned in mice, for which there are many more antibodies and resources available to definitively identify the APCs that process and present the antigens and further elucidate the mechanisms that drive the strong cellular immune responses induced by suction-based vaccination. However, this will require us to adapt the suction device and parameters for treatment of mouse skin, which is much looser than rat or human skin.

## Data Availability Statement

The data of this study is included in the article and supporting information. Further queries can be directed to the corresponding authors.

## Ethics Approval Statement

All animal experiments were approved by Rutgers University Institutional Animal Care and Use Committee (IACUC), protocol: 201800077.

