## Supplemental Information for "Evaluation of Antigen Expression and Early Immune Response following Cutaneous Suction-mediated DNA Delivery"

Y. Um
Department of Animal Sciences, Rutgers University-New Brunswick, New Brunswick, NJ 08901, USA.

S. H. Park
Department of Materials Science and Engineering, Rutgers University-New Brunswick, Piscataway, NJ 08854, USA.

J. D. Zahn, D. I. Shreiber
Department of Biomedical Engineering, Rutgers University-New Brunswick, Piscataway, NJ 08854, USA.

L. K. Denzin
Child Health Institute of New Jersey, Rutgers Robert Wood Johnson Medical School, New Brunswick, NJ 08901, USA.

E. O. Lallow, C. C. Roberts, Y. K. Park, J. N. Maslow
GeneOne Life Science, Inc., Seoul 07335, Republic of Korea.

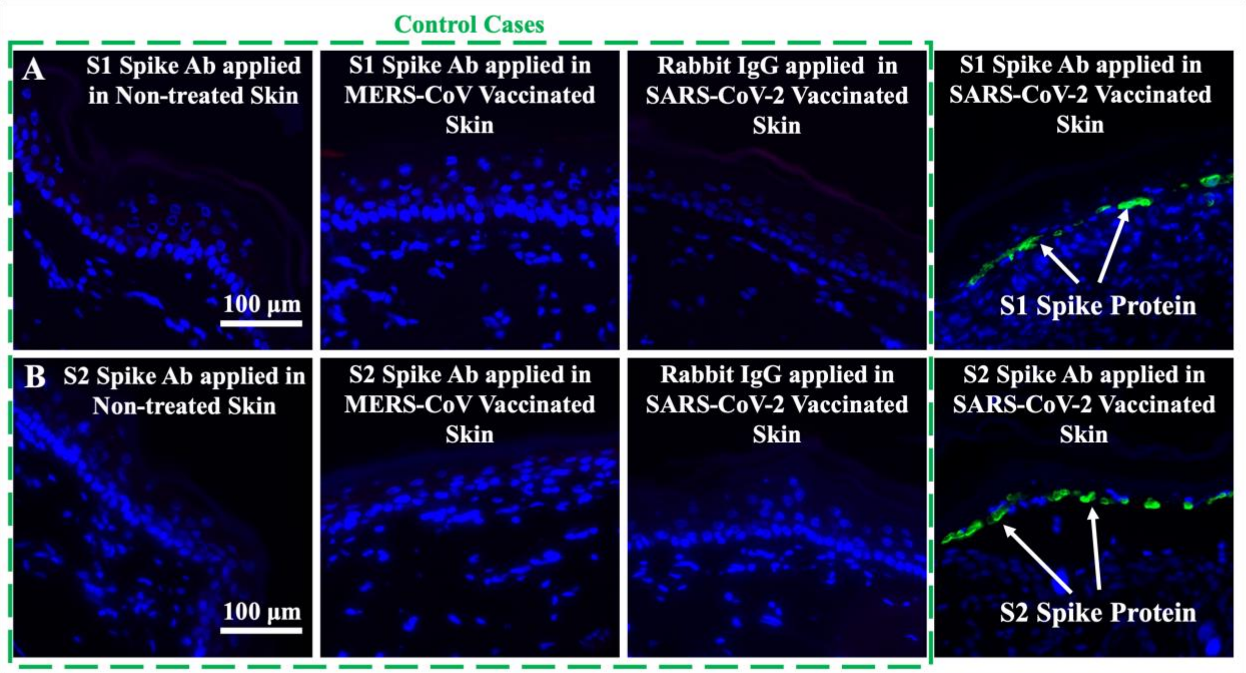


**Figure S1:** Detection of SARS-CoV-2 (A) spike S1 protein and (B) spike S2 protein in rat skin epidermis following suction-mediated vaccination at 24 hours, compared with the control cases: spike S1 and S2 antibody application to non-treated rat skin, and MERS-CoV vaccinated rat skin; and Rabbit IgG application to SARS-CoV-2 vaccinated rat skin.
